# Glass confers rhabdomeric photoreceptor identity in *Drosophila*, but not across all metazoans

**DOI:** 10.1101/446161

**Authors:** F. Javier Bernardo-Garcia, Maryam Syed, Gáspár Jékely, Simon G. Sprecher

## Abstract

Across metazoans, visual systems employ different types of photoreceptor neurons to detect light. These include rhabdomeric PRs, which exist in distantly related phyla and possess an evolutionarily conserved phototransduction cascade. While the development of rhabdomeric PRs has been thoroughly studied in the fruit fly *Drosophila melanogaster*, we still know very little about how they form in other species. To investigate this question, we tested whether the transcription factor Glass, which is crucial for instructing rhabdomeric PR formation in *Drosophila*, may play a similar role in other metazoans. Glass homologues exist throughout the animal kingdom, indicating that this protein evolved prior to the metazoan radiation. Interestingly, our work indicates that *glass* is not expressed in rhabdomeric photoreceptors in the planarian *Schmidtea mediterranea* nor in the annelid *Platynereis dumerilii*. Combined with a comparative analysis of the Glass DNA-binding domain, our data suggest that the fate of rhabdomeric PRs is controlled by Glass-dependent and Glass-independent mechanisms in different animal clades.

## INTRODUCTION

Most animals can detect visual cues, which provide them with detailed information about their environment. This information may include the shape of nearby objects, colours, movements or the day–night cycle, and it is relevant for surviving. As a consequence, animals have evolved various types of photoreceptor neurons (PRs) such as ciliary and rhabdomeric PRs [1, 2], which play different roles in different animal species. For instance, rhabdomeric PRs are critical for image-forming vision (e.g. in *Drosophila* compound eye PRs) and for identifying the direction of a light source (e.g. in the planarian *Schmidtea mediteranea* and in the annelid *Platynereis dumerilii*) [3-5]. Nevertheless, in the case of most metazoan clades, very little is known about how rhabdomeric PRs develop.

Interestingly, all known rhabdomeric PRs appear to use a similar assortment of phototransduction proteins. These PRs possess rhabdomeric-type opsins that can modify their spatial conformation upon light stimulation, which allows them to activate Gαq. Gαq signals through phospholipase C (PLC) causing the opening of cation channels on the cytoplasmic membrane of PRs and thus leading to the formation of action potentials. This light-sensing machinery is present in distantly related animal phyla [1, 6, 7], including vertebrates (due to the ‘intrinsically photosensitive retinal ganglion cells’, ipRGCs [8]), which poses the question of to what degree the development of rhabdomeric PRs is evolutionarily conserved. Is the acquisition of the rhabdomeric phototransduction cascade regulated by a similar set of transcription factors across metazoans? Using the fruit fly *Drosophila melanogaster* as a model system we have recently shown that the zinc finger transcription factor Glass acts as critical cell fate selector by directing the maturation of PR precursors into adult, light-sensing PRs. In *Drosophila*, Glass is required for the expression of virtually all the phototransduction proteins [9], and it regulates the development of all types of rhabdomeric PRs (including those in the Bolwig organ, the ocelli, and the compound eye) [10-12]. Therefore, we investigated whether Glass may also be involved in rhabdomeric PR differentiation in other animal species.

The planarian *Schmidtea* and the annelid *Platynereis* are emerging model organisms whose visual systems have been well characterised [3, 5, 13-19]. Interestingly, by analysing recently published single-cell sequencing data of *Schmidtea* we found that *glass* is not expressed in rhabdomeric PRs in this species. Moreover, using *in situ* hybridisation we could not detect *glass* expression in rhabdomeric PRs in *Platynereis.* Thus, while Glass is critical for the specification of rhabdomeric PR identity in *Drosophila*, the absence of Glass in rhabdomeric PRs in *Schmidtea* and *Platynereis* supports that different genetic programmes are required for controlling rhabdomeric PR cell fate in different animal clades. Therefore, while the initial specification of the eye field appears to be controlled by an evolutionarily conserved group of transcription factors (called the retinal determination network, RDN) [17, 20, 21], the subsequent steps that diversify distinct cell types, including rhabdomeric PRs, are likely controlled by diverse developmental programmes.

## METHODS

### Phylogenetic analysis

We searched for protein sequences similar to *Drosophila* Glass [22] and *Platynereis* Glass [23] (see sequences in Additional file 1 and Additional file 2) by using NCBI BLAST [24] and the *Schmidtea mediterranea* Genome Database [25]. Redundant sequences were removed from the collection using CD-HIT with an identity cutoff of 90% [26]. To obtain cluster maps based on all against all pairwise similarity, we used CLANS2 with the BLOSUM62 matrix and a *p*-value cutoff of 1e-60 [27]. For phylogenetic tree construction we selected a non-derived set of sequences from the *glass* cluster and aligned them with MUSCLE [28]. Sequences shorter than 300 amino acids were removed prior to the alignment. We trimmed the alignments with TrimAI in ‘Automated 1’ mode [29]. We identified the JTT+I+G4 model as the best by IQ-TREE [30]. Maximum likelihood trees and bootstrap analysis were carried out with IQ-TREE. Trees were visualized with FigTree [31] (for the data corresponding to this analysis, see Additional file 3).

### Glass-binding site analysis

We examined a subset of Glass-like protein sequences by aligning them with either BLAST [24] or MUSCLE [28], and analysed them with ‘DNA-binding site predictor for Cys_2_His_2_ Zinc Finger Proteins’ [32, 33] (for details on the sequences that we used see Fig. 3 and Additional file 4). To investigate the DNA-binding specificity of each of these candidates, we copied its full amino acid sequence as input, and asked the software to search for Cys_2_His_2_ domains [32]. Then, we predicted the binding sites for those regions that best aligned with the 4th and the 5th zinc fingers of Glass, which are responsible for recognising its targets *in vivo* [34-37]. We used ‘expanded linear SVM’ as prediction model.

### Animal caretaking

*Drosophila melanogaster* stocks were cultured at 25 °C in a 12:12 hour light–dark cycle, and we fed them with cornmeal medium (which was supplemented with molasses, fructose and yeast). We used Canton-S as a wild-type strain (courtesy of R. Stocker), *glass-Gal4* (courtesy of S. Kim) [38] and *UAS-mCD8::RFP* (Bloomington Stock Center, no. 32219).

Our wild-type *Platynereis dumerilii* were a mixed population of worms, originally captured in the sea in Naples (Italy) and Arcachon (France). We also used *r-opsin1-GFP* worms (courtesy of F. Raible) [14]. These animals were kept in sea water at 22 °C, in a 16:8 hours light–dark cycle. We maintained them synchronised to an artificial moon cycle, induced by slightly increasing the light intensity at night for 1 week per month (using a 10 W light bulb, to simulate the full moon). *Platynereis* had a varied diet that included *Artemia salina, Tetraselmis marina*, fish food and spinach leaves. For our experiments (i.e. *in situ* hybridisation and microinjections) we crossed males and females and collected the fertilised eggs, as previously described [39]. The larvae that hatched from these eggs were kept at 18 °C.

### Immunohistochemistry and in *situ* hybridisation

In the case of *Drosophila* antibody stainings, these were performed on cryosections of *glass>mCD8::RFP* flies, as previously described [9, 40]. We dissected heads (removing the proboscis to improve the penetration of our reagents) and fixed them for 20 minutes with 3.7% formaldehyde in 0.01 M phosphate buffer (PB; pH 7.4). Then, we washed our samples with PBT (Triton X-100 0.3% in PB) and incubated them with a cryoprotectant solution (sucrose 25% in PB) overnight at 4 °C. The following day we embedded the fly heads in OCT, froze them with liquid nitrogen, and cut 14 μm cryosections in the transverse plane. Once the samples were dry, we proceeded to immunostain them. For this, we washed the slides with PBT (this buffer was also used in subsequent washing steps) and incubated them in primary antibody (rabbit anti-DsRed, 1:100, Clontech, no. 632496) at 4 °C overnight. Then, we washed the cryosections and incubated them in secondary antibody (goat anti-rabbit conjugated to Alexa Fluor 568, 1:200, Molecular Probes, no. A-11011) at 4 °C overnight, and washed again the next day. We mounted our samples by using Vectashield that contained DAPI (Vector, H-1200) and took images with a Leica SP5 confocal microscope.

To detect the *glass* transcript in *Drosophila*, we used the ViewRNA *in situ* hybridisation kit of Affimetrix (no. QVT0012) – which is a proprietary method – and proceeded according to the instructions of the company. Briefly, we took head cryosections (as described above for antibody stainings) and ordered a mix of labelled RNA probes against *glass* from Affimetrix. Then, we processed the samples by digesting them with protease QF, and washed with PB and with various commercial solutions included in the kit. We incubated our cryosections with the *glass* probes for 2 hours, at 40 °C. After this, we continued with a series of washing and signal amplification steps, followed by a colour reaction (we used Fast Red as a fluorophore). We finished by washing the samples with PB, and used Vectashield containing DAPI (Vector, H-1200) to cover the slides. Imaging was done with a Leica SP5 confocal microscope.

To perform double *in situ* hybridisation in *Platynereis*, we followed – with few modifications – a protocol that has been previously used for characterising the expression pattern of *r-opsin1* [16, 41]. For the present work, we also produced an RNA probe against the *glass* transcript (for details on the *glass* probe, see Additional file 1). We fixed 3–5 day old larvae in 4% formaldehyde, and we subjected them to a mild proteinase K digestion to improve the penetration of our reagents. These larvae were prehybridised at 65 °C by using a hybridisation mix (Hyb-Mix), containing 50% formamide, 5x saline-sodium citrate buffer (SSC), 50 µg/ml heparin, 0.1% Tween 20, and 5 mg/ml torula RNA. Then, we dissolved the probes against *r-opsin1* and *glass* (labelled with either fluorescein-UTP or digoxigenin-UTP) in Hyb-Mix, denatured them at 80 °C for 10 minutes, and added this solution to our samples. We hybridised both probes simultaneously by incubating at 65 °C overnight. Then, we washed the samples at 65 °C with a series of solutions that initially contained 50% formamide and 2x SSCT (obtained from a stock solution with Tween 20 0.1% in 4x SSC), and we progressively decreased the concentration of both formamide and SSCT throughout successive washes. After washing, we placed the larvae at room temperature and proceeded to immunostain them. We detected the two probes sequentially, by using peroxidase-conjugated primary antibodies against fluorescein (1:250, Roche) and digoxigenin (1:50, Roche). To detect the first probe, we incubated our samples overnight at 4 °C in one of these antibodies, washed them with Tris NaCl Tween 20 buffer (TNT; 0.1 M Tris-HCl, 0.15 M NaCl, 0.1% Tween 20; pH 7.5), and started the colour reaction by adding a solution that contained fluorescent tyramide (conjugated to either Cy3 or fluorescein). We controlled the development of the signal by using a fluorescence microscope and, when it was ready, we washed in TNT and stopped the peroxidase activity with H_2_O_2_. To detect the second probe, we repeated these immunostaining steps similarly. We mounted our samples with 90% glycerol, and scanned them in a confocal microscope (example confocal stacks can be found within the Additional file 5).

### Microinjection of *glass-Tomato*

We used an unpublished assembly of the *Platynereis* genome (courtesy of D. Arendt, EMBL Heidelberg) for making a *glass-Tomato* reporter (see Additional file 1 and Additional file 2 for details). We PCR-amplified a fragment of the *Platynereis glass* promoter and cloned it into a plasmid in frame with the tandem dimer version of *Tomato* (courtesy of L. A. Bezares-Calderón) by using sticky end ligation with ApaI and SgsI [42]. The fragment that we cloned included a 5,789 bp long upstream sequence, and also the beginning of the Glass coding sequence: the first ATG codon was predicted both by aligning the *Platynereis* version of Glass to the Glass homologues of other species and by using the ATGpr software [43, 44]. For details on the plasmid that we injected, see its annotated sequence in Additional file 6.

For microinjections, we collected freshly fertilised *Platynereis* eggs and proceeded as previously described [14]. Briefly, we removed the jelly of the eggs by digesting with proteinase K and washing with abundant sea water, using a sieve. We diluted the *glass-Tomato* plasmid to a final concentration of about 200 ng/μl, and delivered it into 1-cell embryos with a microinjection set-up, by using Femtotip II microcapillaries (Eppendorf). Larvae were kept at 18 °C, and we imaged them with a Leica SP8 confocal microscope to study the expression of the reporter (representative confocal stacks are available in the Additional file 5). The expression of this reporter showed some degree of mosaicism, given that it was not integrated into the genome, which allowed us to investigate the morphology of the individual neurons that expressed it. We investigated over 100 surviving, Tomato-positive *Platynereis* larvae.

## RESULTS

### Glass homologues are present throughout metazoans

Glass plays a fundamental role for the differentiation of rhabdomeric PRs in the fruit fly [9, 11, 45, 46]. To investigate if it provides a similar function across metazoans, we first decided to search for Glass homologues in other species.

To do this, we obtained Glass-like sequences by using NCBI BLAST [24] and the *Schmidtea mediterranea* Genome Database [25]. We analysed these sequences with the CLANS2 software (using the BLOSUM62 matrix and a *p*-value cutoff of 1e-60) to produce a cluster map (Fig. 1A) [27]. In this type of graph, closely related sequences (represented as points) are clustered together and connected by dark lines. Based on their similarities, we were able to identify multiple Glass homologues across distantly related species. Some more derived sequences (e.g. from *Strongylocentrotus* and *Saccoglossus*) were also clearly supported as Glass homologues in our analysis. Using these data, we constructed a maximum likehood phylogenetic tree for Glass, which was visualised with FigTree (Fig. 1B) [31] (for more details on our analysis, see the Methods section and the Additional file 3). Importantly, our data reveal that Glass homologues are widely present throughout the animal kingdom.

**Fig. 1:**
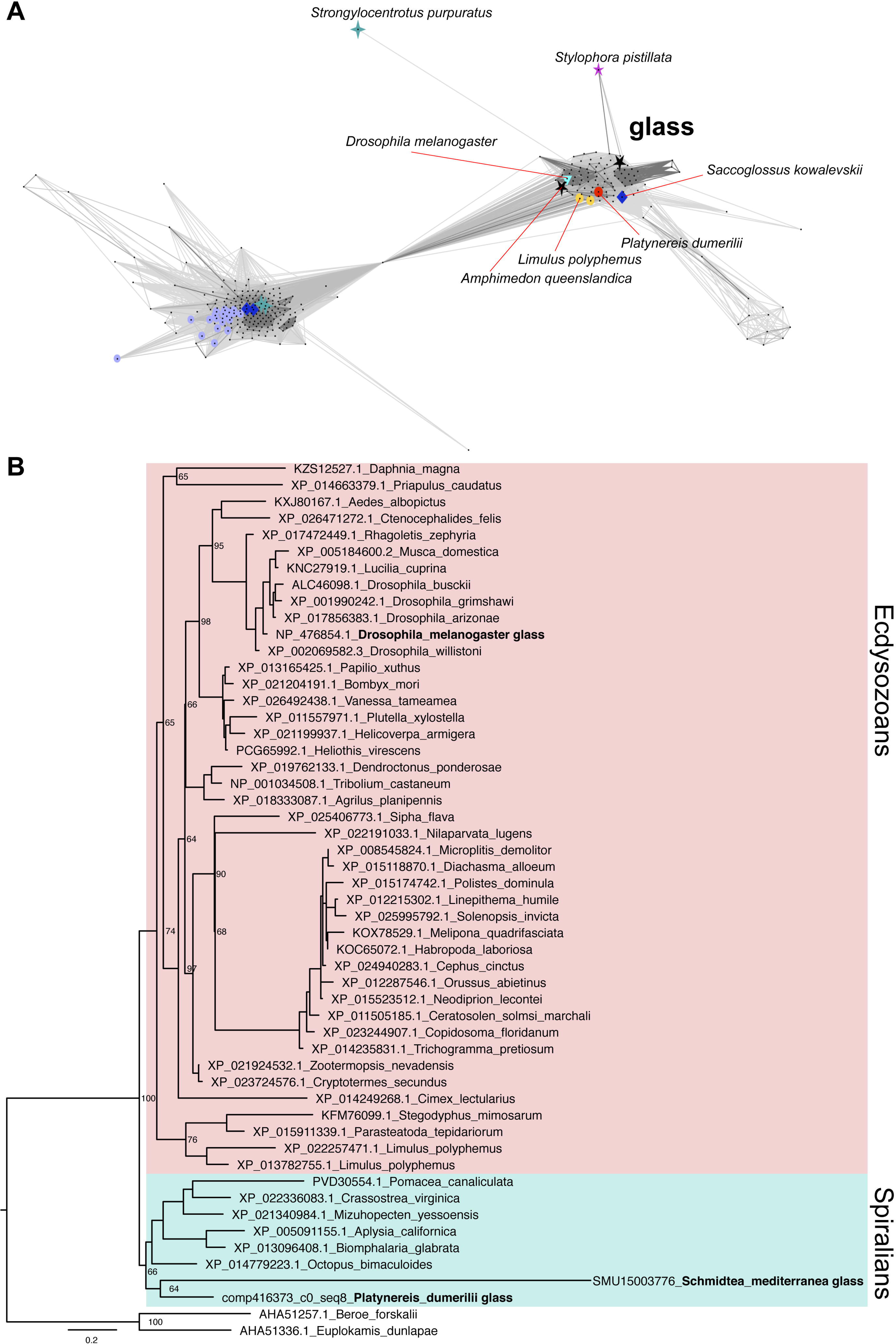
Glass phylogeny. To identify Glass homologues we searched for Glass-like sequences with BLAST and obtained a cluster map by using all against all pairwise similarity. In this graph, those sequences that are most similar appear clustered together, and connected by a darker line (A). Based on these data, we built a maximum likehood tree for Glass (B) (for further details see the Methods section, the tree file and the sequences that we used for it are included in the Additional file 3).

### Neither vertebrates nor choanoflagellates have clear Glass homologues

Based on the distribution of Glass homologues, it seems this protein was present in the common ancestor of all metazoans, but not in choanoflagellates (the sister group of metazoans). Intriguingly, we could also not find any Glass homologue in vertebrates (Fig. 2). Since we identified Glass across multiple animal clades, we wondered why we could not find its vertebrate homologue. Several species have fully sequenced, well annotated genomes, like zebrafish, mice, or humans [47-51]. For this reason, we decided to further investigate the evolutionary conservation of Glass by scrutinising its sequence.

**Fig. 2:**
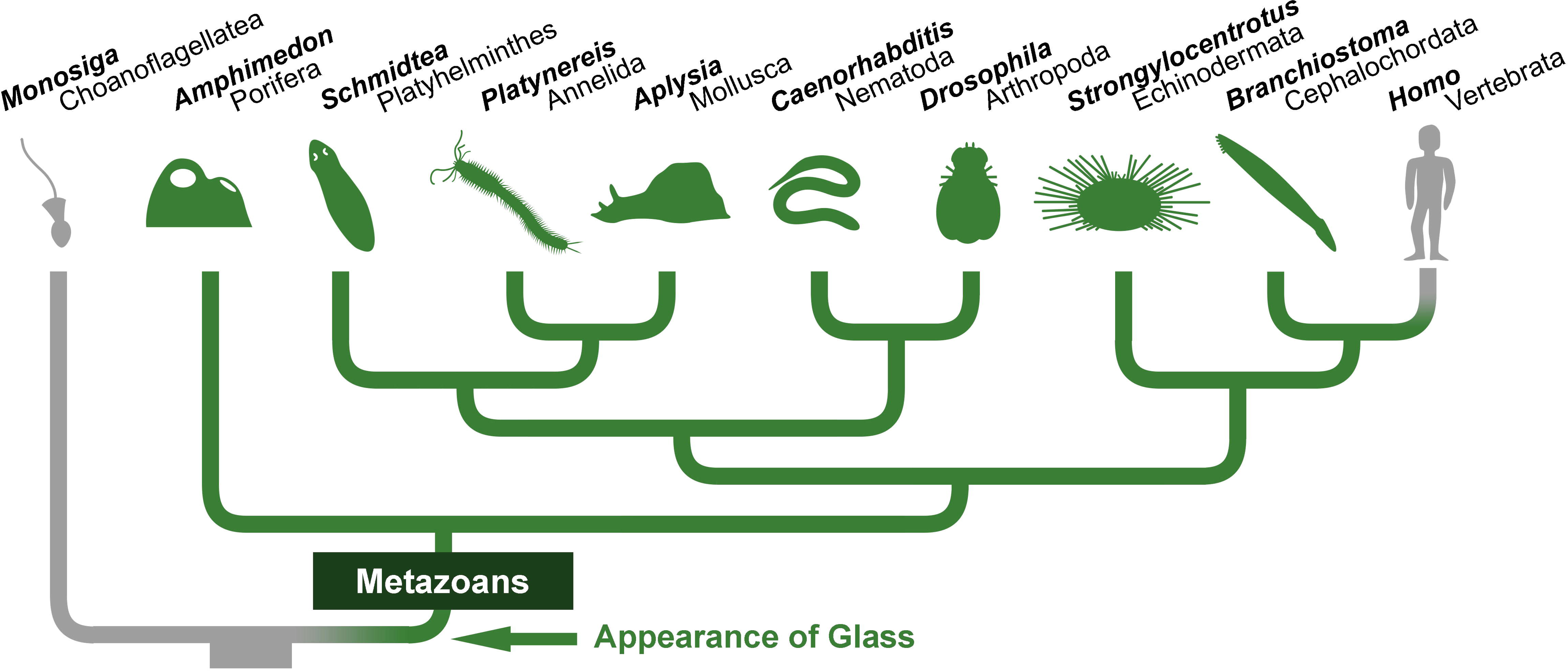
Glass homologues exist in most animal groups. Based on sequence comparison (Additional file 4, also see Fig. 3), we infer that *glass* appeared in the common ancestor of all metazoans, and that it has been transmitted to most present-day animals (shown in green on the phylogenetic tree [74]). However, we were not able to identify *glass* in vertebrates.

**Fig. 3:**
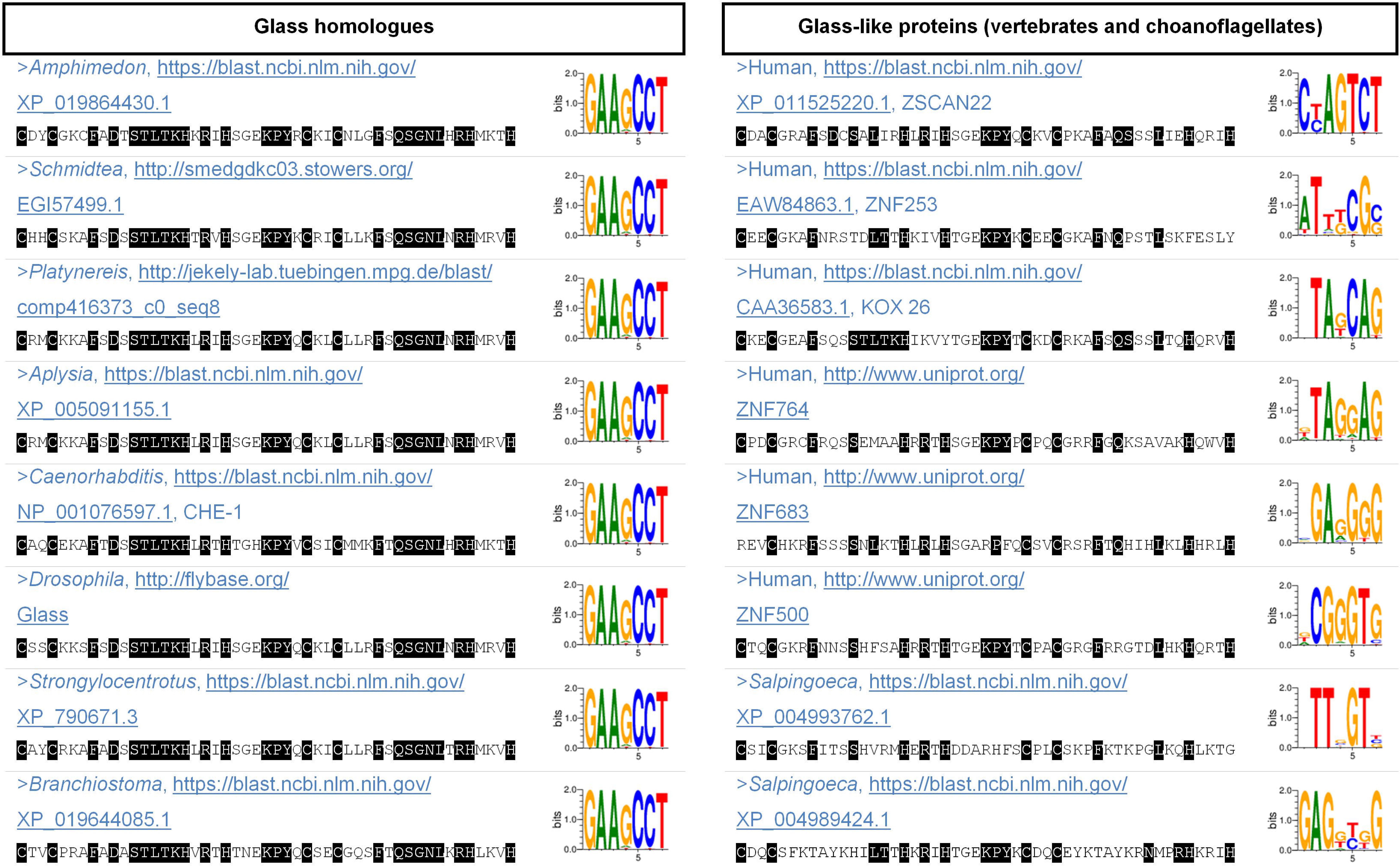
Analysis of the Glass zinc fingers. Generally, Glass homologues possess a cluster of five Cys_2_His_2_ zinc fingers, each of them containing the following motif: Cys-X_2,4_-Cys-X_12_-His-X_3,4,5_-His. Of these, we compared the sequences of the 4th and the 5th zinc fingers, which are responsible for recognising the DNA Glass-binding motif in PRs *in vivo* [34-37], from the following species: *Amphimedon* (Porifera), *Schmidtea* (Platyhelminthes), *Platynereis* (Annelida), *Aplysia* (Mollusca), *Caenorhabditis* (Nematoda), *Drosophila* (Arthropoda), *Strongylocentrotus* (Echinodermata) and *Branchiostoma* (Cephalochordata). In the table, those amino acids that match the Glass consensus sequence (deduced by aligning the homologues of different species, in the first column) appear on black background. The 3D structure of the DNA-bound Cys_2_His_2_ domain has been resolved [75], and it is expected that four amino acids per zinc finger directly recognise three base pairs. These amino acids are well evolutionarily conserved across different Glass homologues and, in the sequences that we show, they are no. 10 (D), 12 (S), 13 (T), and 16 (K) in the 4th zinc finger, and no. 38 (Q), 40 (G), 41 (N), and 44 (R) in the 5th zinc finger. Other residues and neighbouring zinc fingers are also expected to contribute to the DNA-binding specificity of Glass [76]. Similarly, we aligned Glass-like proteins from vertebrates (e.g. human) and choanoflagellates (e.g. *Salpingoeca*) with BLAST [24] and MUSCLE [28], but they showed little similarity to the Glass consensus sequence (shown on the second column). Furthermore, a ‘DNA-binding site predictor for Cys_2_His_2_ Zinc Finger Proteins’ has been developed and is available online [32, 33]. This software predicts that, based on their amino acid sequence, all Glass homologues (in the first column) can bind to the same DNA motif:GAAGCC, which was expected from experimental works in *Drosophila* and *Caenorhabditis* [34, 35]. By contrast, it appears that the Glass-like proteins of vertebrates and choanoflagellates (on the second column) would not be able to recognise this motif. All sequences are available in the Additional file 4.

Glass homologues share a distinctive cluster of five Cys_2_His_2_ zinc fingers in most species (one exception is *Caenorhabditis*, in which it only has four zinc fingers because the first one is missing). Particularly, the 4th and the 5th zinc fingers are especially important because they are responsible for guiding Glass towards its targets, given that they recognise its DNA-binding motif *in vivo*, GAARCC [34-37]. Therefore, we modified our bait by using the consensus sequence of either the full cluster of five zinc fingers, or only the 4th and 5th zinc fingers, and we repeated our BLAST search against vertebrates and choanoflagellates. By doing this we obtained results like, for example, ZSCAN22, ZNF253, or KOX 26 in humans, which still showed less similarity to Glass than any of those homologues that we identified in other species (Fig. 3, sequences available in Additional file 4). We also considered the human candidates that appeared annotated as putative Glass orthologues in Flybase via DIOPT [22, 52], including ZNF764, ZNF683, or ZNF500, but, likewise, they aligned poorly with the consensus sequence of the Glass zinc fingers (Fig. 3, sequences available in Additional file 4). Next, we analysed if any of these proteins would be able to functionally substitute Glass by recognising its DNA-binding motif, the GAARCC sequence [34, 35, 37]. For this, we used the online tool ‘DNA-binding site predictor for Cys_2_His_2_ Zinc Finger Proteins’, which predicts the DNA-binding behaviour of zinc finger proteins [32, 33]. This software indicates that those Glass-like proteins that exist in vertebrates and choanoflagellates cannot recognise the GAARCC motif, in contrast to the clear Glass homologues that we found in other animals (i.e. in *Amphimedon, Schmidtea, Platynereis, Aplysia, Caenorhabditis, Drosophila, Strongylocentrotus* and *Branchiostoma*) (Fig. 3). Consequently, it remains unclear what happened to the *glass* gene during the evolution of vertebrates: it could be that they lost Glass, or that it severely changed its amino acid sequence and its DNA-binding motif. Intriguingly, similar to *Drosophila*, some cells in the vertebrate retina also use the rhabdomeric phototransduction cascade – the ipRGCs, which detect irradiance [8] – and, based on our data, it seems highly probable that these cells develop through different mechanisms in *Drosophila* and in vertebrates.

### *glass* is not expressed in rhabdomeric PRs in the *Schmidtea* eye

Given that Glass is an essential transcription factor for activating the expression of phototransduction proteins in all *Drosophila* PRs [9, 10], we investigated whether Glass has a similar function in other organisms. For this, we tested if it is expressed in PRs in the eye of the planarian *Schmidtea mediterranea*. Planarians typically possess one pair of eyes, located in the head, that mediate light avoidance [5, 17, 53]. Importantly, their eyes contain rhabdomeric PRs, which are evolutionarily homologous to *Drosophila* PRs [1, 17].

Recently, a single-cell transcriptome atlas has been published for *Schmidtea*, and it is available online [18, 19, 54]. Using this database, rhabdomeric PRs can be identified because they form a cluster of non-ciliated neurons that express phototransduction proteins, including the *opsin* gene (Fig. 4A) [19]. Surprisingly, these cells do not co-express Glass (Fig. 4B), suggesting that, in contrast to *Drosophila*, Glass is not important for the function of rhabdomeric PRs in the *Schmidtea* eye.

**Fig. 4:**
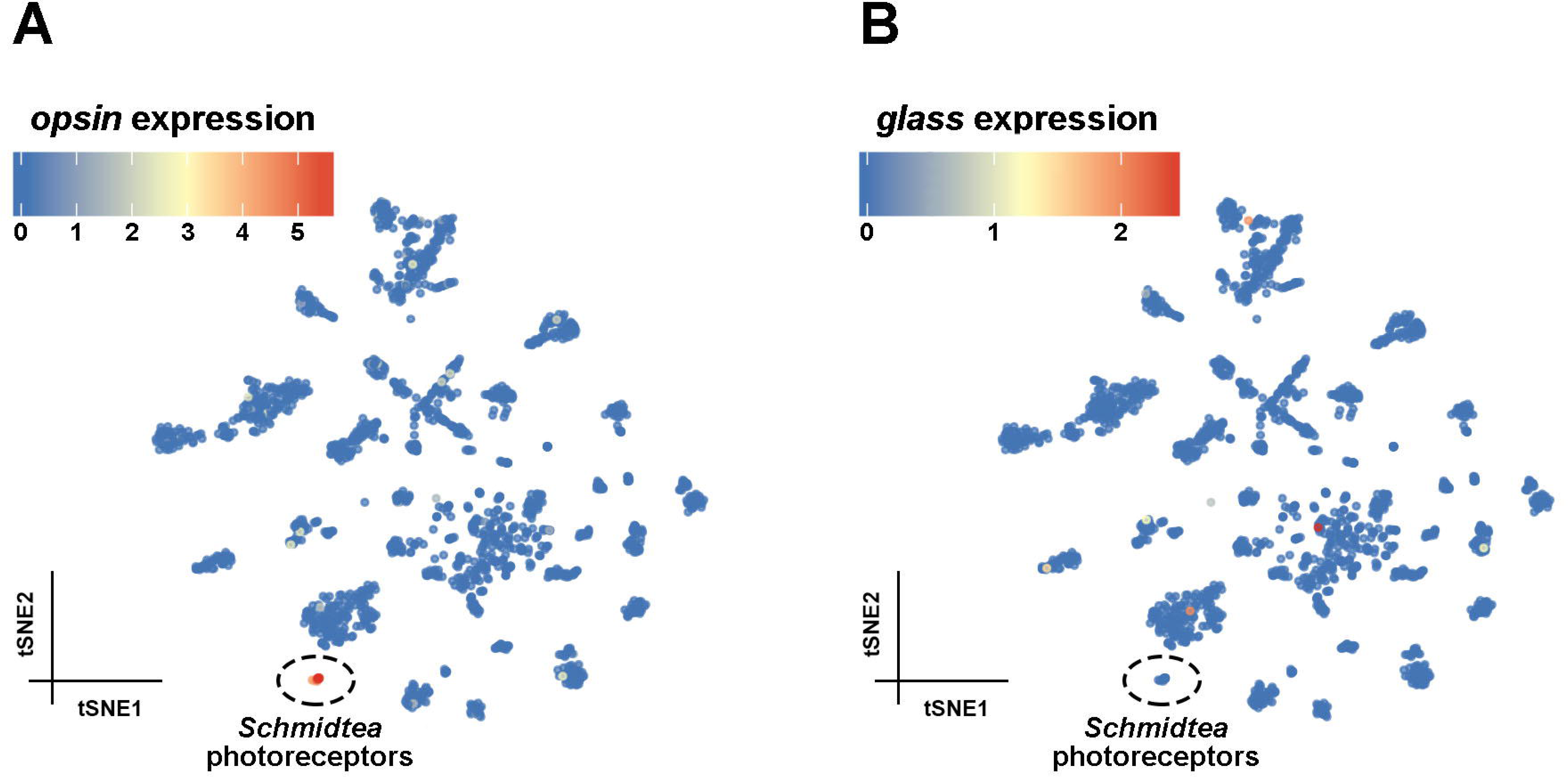
*glass* is not expressed in rhabdomeric PRs in *Schmidtea*. These graphs were obtained from the Planarian Digiworm atlas, a single-cell transcriptome database for *Schmidtea mediterranea* [19, 25]. Each point corresponds to one single cell, and they are clustered according to the similarity of their transcriptome. One of the clusters shown, corresponding to non-ciliated neurons, is formed by 14 rhabdomeric PRs which can be identified because of the expression of the *opsin* gene (dd_Smed_v4_15036_0_1, A). However, these PRs do not appear to express the *Schmidtea glass* homologue (annotated as dd_Smed_v4_75162_0_1 in this website [19, 54], B).

### *glass* is not expressed in rhabdomeric PRs in the *Platynereis* eye

We next tested whether Glass is expressed in rhabdomeric PRs in the marine ragworm *Platynereis dumerilii*. The visual system of *Platynereis* has been well studied, both from a molecular and a functional point of view. *Platynereis* possesses two types of bilateral eyes containing rhabdomeric PRs, called the dorsal and ventral eyes (also known as adult and larval eyes, respectively). These two eye types are able to detect the direction of light, thus mediating phototaxis [3, 13-16].

In *Drosophila, glass* is expressed in all rhabdomeric PRs [12, 55]. We could detect *glass* expression in the compound eye of adult flies both with *in situ* hybridisation and with a *glass-Gal4* line crossed to *UAS-mCD8::RFP* (Figs. 5A–B′), which confirms previous data [12, 55]. By contrast, in the case of *Platynereis, in situ* hybridisations performed in 3–5 day old larvae did not show co-expression of the *glass* transcript with *rhabdomeric opsin 1* (*r-opsin1*), which is a marker for rhabdomeric PRs in both the dorsal and the ventral eyes [14, 16], indicating that *glass* is not present in these cells (Figs. 5C–C′′′′, also see confocal stacks in Additional file 5). In addition, we also generated a *Platynereis glass* reporter by cloning 5.7 kb of its upstream sequence into a plasmid, where the *glass* start codon was in frame with *Tomato* (a red fluorescent protein). We used this plasmid for transient transgenesis by injecting it in 1-cell embryos containing a stable *r-opsin1-GFP* insertion [14]. *r-opsin1-GFP* animals consistently showed strong GFP signal in their dorsal eye PRs, and this signal was weaker in the ventral eye PRs. In the case of the dorsal eyes, all PRs project their rhabdomeres into a pigment cup, and their axons form four nerves that innervate the optic neuropil in the brain [3, 14, 16]. After microinjections, we tested 3–8 day old larvae (slightly older than those that we used for *in situ*, to guarantee that positive cells had enough fluorescence to distinguish them) but we did not observe co-expression of GFP and Tomato. *glass-Tomato*-expressing neurons were consistently located in the head of *Platynereis*, distant from the ventral eyes. The expression of *glass-Tomato* showed some degree of mosaicism due to this reporter not being integrated into the genome, which allowed us to observe the morphology of individually labelled cells in detail. Some of these Tomato-positive cells appeared close to the dorsal eyes, but they did not project any rhabdomere-like extension into the pigment cup, and their axons did not innervate the optic neuropil (Figs. 5D–E′ ′, confocal stacks are available in Additional file 5), indicating that they were not part of the eye rhabdomeric PRs. We conclude that, while Glass is expressed in all types of rhabdomeric PRs in *Drosophila*, it is not present in known rhabdomeric PRs in *Platynereis*.

**Fig. 5:**
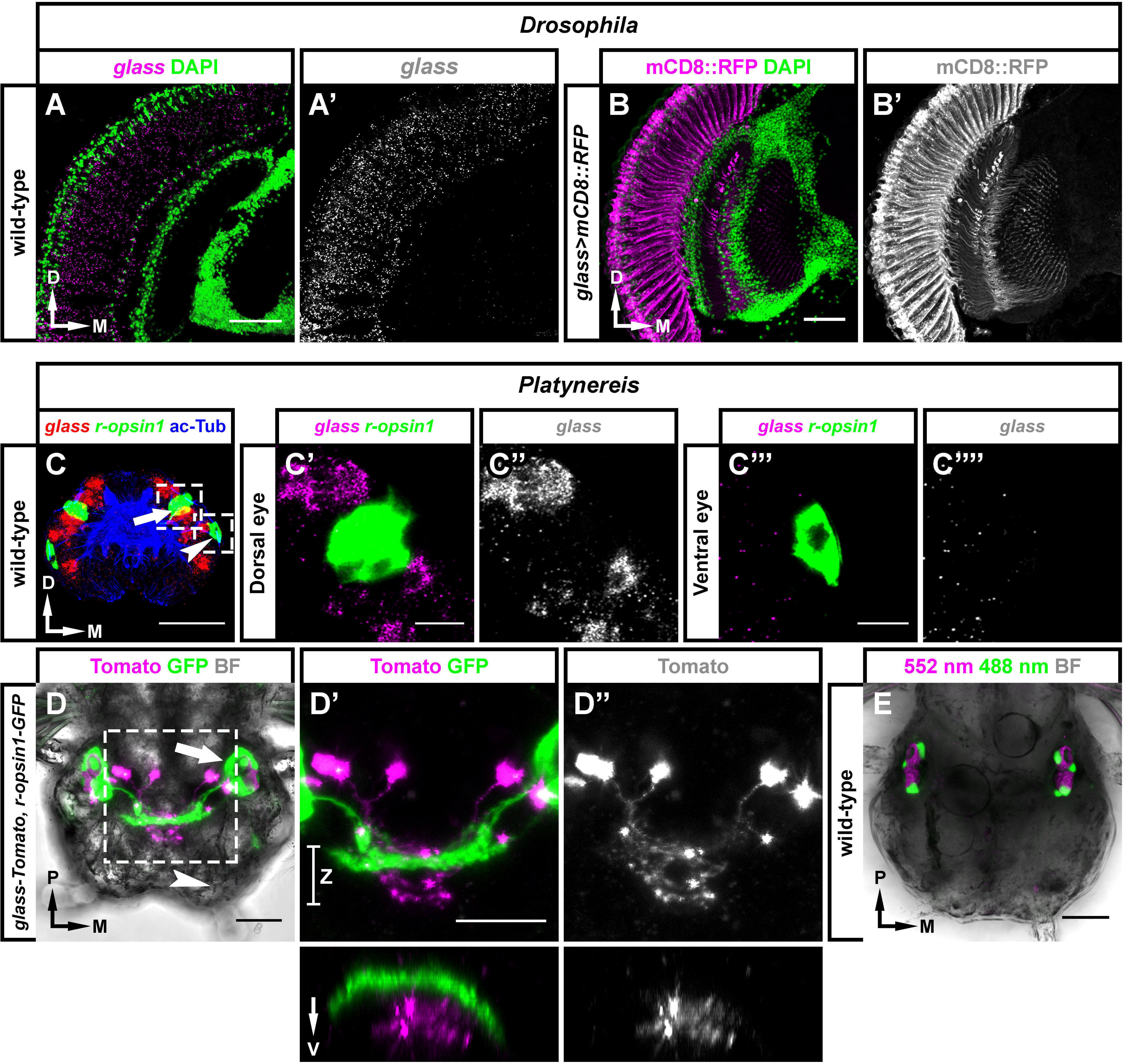
*glass* is not expressed in rhabdomeric PRs in *Platynereis*. (A, B) *glass* is present in all *Drosophila* rhabdomeric PRs, including those in the compound eye [12, 55]. This can be observed in head cryosections, either by using *in situ* hybridisation (magenta in A, greyscale in A′) or with *glass>mCD8::RFP* flies (magenta in B, greyscale in B′). In both cases samples were counterstained with DAPI (green). (C-E) In contrast to *Drosophila*, double *in situ* hybridisation against the *glass* (red) and *r-opsin1* (green) transcripts shows that *glass* is not present in *Platynereis* rhabdomeric PRs. Samples were counterstained with antibodies against acetylated Tubulin (ac-Tub, blue), which is a neuropil marker (C, transversal view of a whole-mounted, 5 day old larva). To the right, close-ups of the dorsal (arrow in C; C′,C′′) and ventral eyes (arrowhead in C; C′′′, C′′′′) show that *glass* (in magenta/greyscale) is not expressed in either of these visual organs. Similarly, we found that a microinjected *glass-Tomato* reporter (magenta/greyscale) was not co-expressed with a stable *r-opsin1-GFP* insertion (green). Brightfield (BF, greyscale) was imaged as a reference (D–D′′, dorsal view of a whole mounted, 8 day old larva). The positions of the dorsal and ventral eyes are shown with an arrow and an arrowhead, respectively. Close-ups to the right show how the axons of Tomato and GFP-positive neurons project into two different areas in the brain (D′, D′′; orthogonal views taken along the Z segment are shown below). As a control, we also imaged an 8 day old, wild-type, uninjected larva to test its autofluorescence (using two excitation laser wavelengths: 552 nm, same as for Tomato; and 488 nm, same as for GFP). Scale bars: 10 μm in C′, C′′′; 20 μm in D– E; and 50 μm in A, B. Axes: D, dorsal; M, medial; P, posterior; V, ventral.

### Glass is expressed in *Platynereis* sensory neurons

Since *glass* is predominantly expressed in PRs in *Drosophila*, we wondered what type of cells express *glass* in *Platynereis*. We observed that most of the neurons that were labelled with the *glass-Tomato* reporter innervated the neurosecretory neuropil (which is ventral to the optic neuropil, Figs. 5D–D′ ′) [56], and, interestingly, many of them were bipolar neurons (Fig. 6). These two features are relevant because an ongoing electron microscopy (EM) connectome reconstruction shows that, in *Platynereis* larvae, most neurons are either unipolar or pseudounipolar, and there are very few bipolar neurons [3, 56-59]. Based on their position and on their morphology, all bipolar neurons in this EM reconstruction are considered sensory neurons because they possess distinctive membranous specialisations (called sensory dendrites) that project towards the surface [3, 56-59]. Therefore, it is very likely that a subset of *glass*-expressing cells in *Platynereis* are sensory neurons.

**Fig. 6:**
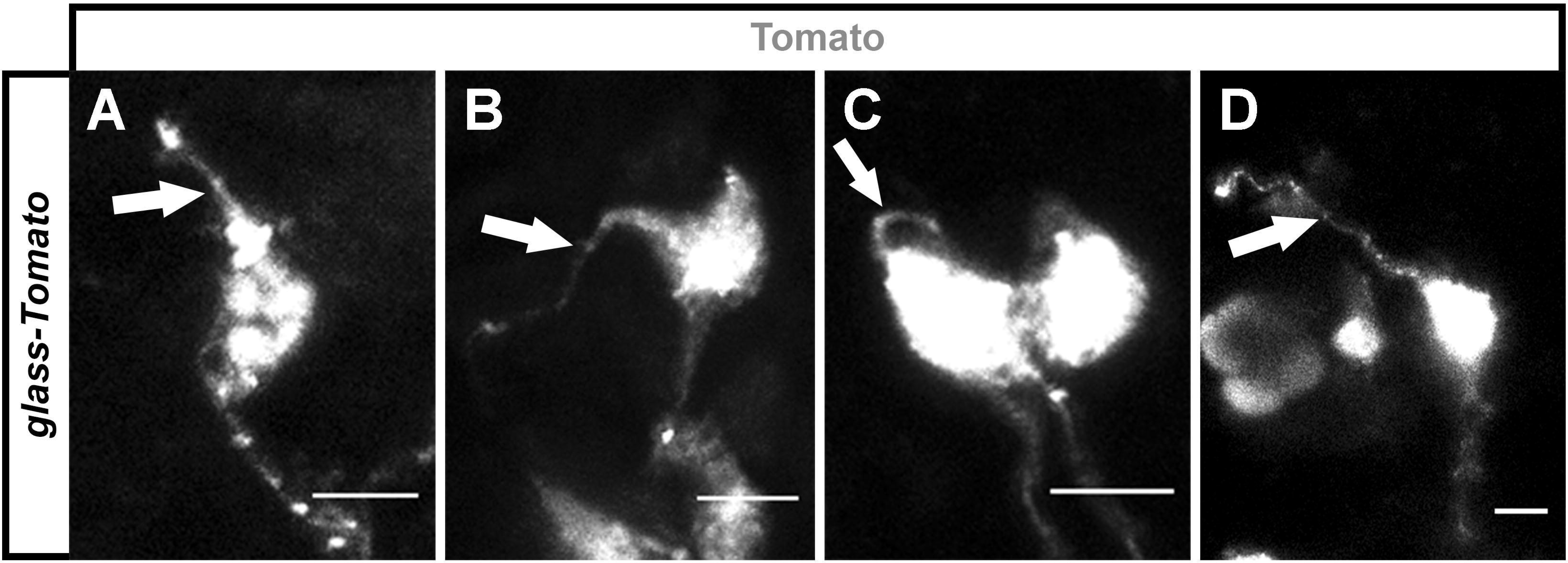
Glass-expressing cells in *Platynereis* include sensory neurons. When we injected our *glass-Tomato* reporter, we observed that many of the neurons that appeared labelled in the *Platynereis* head were bipolar, located close to the surface, and they often possessed membranous specialisations resembling sensory dendrites (arrows) (A–D). Scale bars: 5 μm.

The neurosecretory neuropile of *Platynereis* contains multiple sensory neurons, and it has been characterised both from an anatomical and a molecular point of view [56]. However, it is still unknown whether this region is homologous to any structure of the *Drosophila* brain. Given that *glass* is also required for the development of the corpora cardiaca in *Drosophila* [60], it could be possible that Glass has an evolutionarily conserved function in neurosecretoy cells. In addition, it could also be that Glass regulates the formation of other sensory neurons. Notably, the *Caenorhabditis* homologue of Glass (called CHE-1) is expressed in ASE chemosensory neurons, and it regulates their development [34, 61].

## CONCLUSIONS

Remarkably, the earliest steps of eye development are controlled by a group of transcription factors, called the ‘retinal determination network’ (RDN), which is both required and sufficient for eye formation in distantly related species [20, 62-68]. RDN members, such as Eyeless, Sine oculis, or Eyes absent are important for inducing eye field specification. To achieve this, they establish complex epistatic interactions with each other. These interactions occur similarly across model organisms, suggesting that this is an evolutionarily conserved process [20, 69]. In contrast to the early steps of eye field specification, subsequent mechanisms that specify the cell fate of PRs are not well understood. Here we provide evidence that, during the late stages of eye development, rhabdomeric PRs mature through different mechanisms in different species.

In *Drosophila*, we have recently shown that Sine oculis (a core component of the RDN) directly activates the expression of the transcription factor *glass*, which is crucial for activating the expression of virtually all the phototransduction proteins in all types of *Drosophila* PRs [9, 10, 70]. Based on similarities in their light-sensing machinery, *Drosophila* PRs are considered homologous to the ipRGCs of vertebrates, and also to the rhabdomeric PRs that exist in *Schmidtea* and *Platynereis* [1, 6, 7, 15, 17, 19]. Intriguingly, while we did identify Glass homologues in most metazoans, we could not find a clear Glass homologue in vertebrates. Moreover, our data indicate that *glass* is not expressed in the rhabdomeric PRs of *Schmidtea* or *Platynereis*. This suggests that metazoans have evolved alternative transcriptional pathways to direct the formation of rhabdomeric PRs. One of these pathways requires Glass (e.g. in *Drosophila*), while others do not, (e.g. in vertebrates, *Schmidtea*, or *Platynereis* larvae).

It could be possible that Glass started being expressed in rhabdomeric PRs at some point during the evolution of ecdysozoans and that it became specialised in regulating the differentiation of these cells. Therefore, comparing the differentiation of Glass-expressing and non-Glass-expressing PRs provides a valuable entry point to dissect shared and dissimilar aspects of the developmental programme. Additionally, it would also be interesting to know the identity of Glass-expressing cells for understanding the ancestral function of Glass. The *glass* transcript is rare and lowly expressed in the *Schmidtea* single cell transcriptome data that we have currently available [18, 19], and it was also lowly expressed in the single cell transcriptome datasets of *Platynereis*, to the point of being removed from the analyses of the two papers in which the sequencing was published [71, 72], which makes it impossible to compare the function of *glass*-expressing cells between different species at this moment. It could be possible that this is because only a few cells in the brain express Glass, and these may not have been included in the samples that were sequenced. Therefore we expect that, in the near future, increasing both the number and the quality such single cell transcriptomes for these and other species will be useful to address several questions about the evolution of specific cell fates. For example, some opsins may have other functions apart from light-sensing [73], and it would be relevant to know if *glass* regulates the expression of any such opsin outside the *Platynereis* eye (for example), at any stage.

The absence of Glass in rhabdomeric PRs in the eye of some species argues for other transcription factors being capable of activating the expression of phototransduction proteins, however the underlying mechanism remains unknown. Our data support a rather complex scenario for the evolution of rhabdomeric PRs, but future works on the targets of the RDN may help to better understand how rhabdomeric PR identity is regulated.

## ABBREVIATIONS

ac-Tub: acetylated Tubulin
EM: electron microscopy
PB: phosphate buffer
PBT: phosphate buffer with Triton X-100
PR: photoreceptor neuron
RDN: retinal determination network
r-opsin1: rhabdomeric opsin 1
SSC: saline-sodium citrate buffer
SSCT: saline-sodium citrate buffer with Tween 20.

## DECLARATIONS

### Ethics approval and consent to participate

Not applicable.

### Consent for publication

Not applicable.

### Availability of data and material

All data presented in this article are accessible online, as indicated in either the main text or in the Additional file section.

### Authors’ contributions

F.J.B.-G., G.J., and S.G.S. conceived the study. F.J.B.-G., G.J. and M.S. performed the experiments. F.J.B.-G., G.J., and S.G.S. wrote the manuscript.

#### Acknowledgements

We thank the Bloomington Stock Center, R. Stocker, and S. Kim for fly stocks; F. Raible for the *r-opsin1-GFP Platynereis* strain; D. Arendt for providing access to the draft *Platynereis* genome; and L. A. Bezares-Calderón for plasmids. We are also grateful to C. Adler, J. Cleland, and to our colleagues of the Sprecher and Jékely labs for valuable discussions.

## ADDITIONAL FILES

**Additional file 1:** supplementary methods.

**Additional file 2:** *Platynereis glass* supplementary nucleotide sequences (both genomic and transcriptomic).

**Additional file 3:** Glass phylogenetic tree and sequences data.

**Additional file 4:** subset of *glass*-like sequences for which the DNA-binding affinity was investigated.

**Additional file 5:** example confocal stacks.

**Additional file 6:** annotated sequence of the *glass-tomato* plasmid that was used for

*Platynereis* microinjections.

