## Supplementary material for "Glass confers rhabdomeric photoreceptor identity in *Drosophila*, but not across all metazoans": Suppl 1

### ADDITIONAL FILE 1: SUPPLEMENTARY METHODS

We found a Glass homologue in the *Platynereis* transcriptome database (http://jekely-lab.tuebingen.mpg.de/blast/) and we compared this sequence with of an unpublished reconstruction of the *Platynereis* genome (courtesy of D. Arendt, see Additional file 2). Both sources indicated that *Platynereis* possesses one single Glass homologue, and that two different Glass isoforms can be produced by alternative splicing. These are:

>Glass isoform 1, *Platynereis dumerilii*
MLNVRMTVDVPLCAKNTSYQKPQARMESCYLSAGGHSHGGHHGHGSHGGGGPGGHCGGGGGGSPGPSYYT
SSAAAAVAAAGELWRSSPLKPGSPASASEVCGPPRSSVDLSVNTFAMPPLDIDPLSNFFSFSSPAYKELS
VFKEKGPQDIADALLSLKHAVVHPGMNGQLSPLSPGLPPHLPQSISSAMAPSSSLSSYPMSHQHSQLQSY
GMSSQYGSPPPPPPPAPQYGETGSGQASPCHPQAPQHHSMFPAMSVNVSMNMNVAMGNQYNNSMDNWGHH
PTHQPSAQYSPAAAAAAAQMTSQYPSYGHHHHHHHHHHQNHSYSASYAFSPELRSSSSVSVRDMMYPHPH
KPSSASDSYKDVSSKLYSLSAMRRSPRGSCSPVGLGPPPGLRSHHPSGHLGSASSVDSKVNLCRICGKTY
ARPSTLKTHMRTHSGEKPYRCQTCSKSFSQAANLTAHLRTHSGEKPFRCPMCDRRFSQSSSVTTHMRTHS
GERPYRCRMCKKAFSDSSTLTKHLRIHSGEKPYQCKLCLLRFSQSGNLNRHMRVHANNA

>Glass isoform 2, *Platynereis dumerilii*
MESCYLSAGGHSHGGHHGHGSHGGGGPGGHCGGGGGGSPGPSYYTSSAAAAVAAAGELWRSSPLKPGSPA
SASEVCGPPRSSVDLSVNTFAMPPLDIDPLSNFFSFSSPAYKELSVFKEKGPQDIADALLSLKHAVVHPG
MNGQLSPLSPGLPPHLPQSISSAMAPSSSLSSYPMSHQHSQLQSYGMSSQYGSPPPPPPPAPQYGETGSG
QASPCHPQAPQHHSMFPAMSVNVSMNMNVAMGNQYNNSMDNWGHHPTHQPSAQYSPAAAAAAAQMTSQYP
SYGHHHHHHHHHHQNHSYSASYAFSPELRSSSSVSVRDMMYPHPHKPSSASDSYKDVSSKLYSLSAMRRS
PRGSCSPVGLGPPPGLRSHHPSGHLGSASSVDSKVNLCRICGKTYARPSTLKTHMRTHSGEKPYRCQTCS
KSFSQAANLTAHLRTHSGEKPFRCPMCDRRFSQSSSVTTHMRTHSGERPYRCRMCKKAFSDSSTLTKHLR
IHSGEKPYQCKLCLLRFSQSGNLNRHMRVHANNA

To do *in situ* hybridisation against Glass, we obtained a plasmid from a *Platynereis* EST library, and we generated an RNA probe by using T7 RNA polymerase. We checked the size of our probe by running it in an agarose gel next to an RNA ladder. The plasmid contained the following sequence (uploaded to GenBank, with accession number MK343127):

>DNA template for *glass* RNA probe
CTTGCTCCAAATCTTTCTCTCAAGCTGCTAATCTCACTGCACACTTGCGCACCCACTCTGGCGAGAAACC
TTTCAGATGTCCGATGTGTGACCGCCGTTTTTCCCAGTCGTCTTCGGTGACCACTCACATGAGAACACAT
AGCGGAGAACGGCCTTACAGATGTCGGATGTGCAAGAAGGCCTTCTCCGACAGCTCAACCCTCACCAAGC
ACCTAAGAATCCACAGTGGCGAAAAACCTTACCAGTGCAAACTCTGTTTATTAAGATTTTCTCAGTCTGG
AAATCTAAATAGGCACATGCGAGTACATGCTAACAATGCCTGATTATATTCTGAGCTATTCTGCTACTTC
ATTCATTCTGTGGACTGTGAACAATACCATAATGATCTGACAAGTTGTGACAGTTGTGAATTTACTGCTG
TAGTCTGTGGAATACAGCAATGTGAACTATGGACAGTTTCTTTTTAAATGTTGACAATGGACAATGTCTA
TTATGACTGTAACATTCGTGACATGAATGTGTTGTTTAATGATCAAAACATGTCATAGTTAATATATTTC
TTCATGACAACAAAATCTTGGTAATACACTTTCTTCTTTCCGCCTTGAAATTATTGTAGATTCAATAGTT
ATACCAATCCAATTGTTTTCATAATGATTTTTTCCCTTGCCTGTTGTTTTTTCAAATAAATTCCTTGATC
AATTCCTTCTATTTCTTTTTTACTTCCTTCACTTCTTTTTTTTTCTCCATTTCGCTTTCCTTCCATAGGCC
TAATCCTTCTTTCCTTGCTTCCTTCATTGCCTGATTCACATTCCTTCTGTCTTCCTTCTTGTTGCATCCT
CTCCTTGCTTCTTTGTCTATCACTCATTCCTTTCTTCCTCGCTTCTTTCTATTTTTCCTTCCTTCCTTCC
TTTGCTTTCTCGTAACAATTGCTTGCTTGCTTCATTACTTTCTTCCTTCACCACTTGCTTCCTTCCTTGC
TCACTTTCTTGCCTCCTTCCTTTCTTGTTTCCTTCTCTTCTTCCTTCTGTGCTTCCTATTCTGCTTCATT
CCTTGCTTCCTACCTTGTTTGATTCCATAATCTTTTTCCTTCCTTGCTTTCTTGATAGTTTCTTCCTTGC
TTGCTTTCTTGAAAGTGTCTTGCTTCCTTGATTCATAAAAACTATAAAACAAATTTATACATACACACAA
CTTCAGATTCTTTTTGCTACACATGATTTGAATGTGAAACCATTTGTGCATAATATGTTGATATACATTT
GTTCTACTGGTGCTATTGACTATGATGTTTTATTCTGTCTTTACACTTAATTTCTTTACATCTCTTCTTT

ATTGATATGTAAAATTGTTCATTGCTGTTATTTTTGTTTTGTATATATAATAAACATTTTGTCTTGACTG
TGAAAAAAAAAAAAAAAAAAAAAAAAAAAAAAAAAAAAAAAAAAAAAAAAAAAAAAAAAAAAAAAAAAAA
AAAAAAAAAAAAAAAAAAAAA

To generate a *glass-Tomato* reporter (for the full sequence and plasmid map, see Additional file 4), we amplified the following sequence of the *glass* promoter by using *Platynereis* genomic DNA as a template (ApaI and SgsI were used as restriction enzymes, cut sites are shown in lowercase letters, primer sequences are underlined, start codon appears in green and italics):

>Fragment of the *glass* promoter
aagggcccGCATCCGTGCTTGACAGAATGTGGAGTCAGGTGTTTGCACATCTTCATCACAATACAACAGC
TCCTATAAGGCTCATCCAATGGAAGGTCATGATGCCAAGTTTTGCATAAAAGGTTCTAGCCTGTTCCACT
GACGGTTATTGCCTGCATGGGTTCCAACCCGGAATGGGGAACCCACAATTTACAGCCACTGTGGACCTTC
ATGAGAAAAAATCCAGCTGGATGTCGTTCGGATAAGATGTTTATAATTGGAGGTCCCTTGCATCAGTGAC
TTTCAGTTTCTGTGCTGGGCAAGTAAAATATCCCTCACATGGCGATGATGGCGTAATATAAATGTGTACC
GGTAATGCCTAGATCACACGAGAGTGGTTAAGTGACGGCGAAGTTGTAGAGGTGCCAGGAACTCAAGTTT
CGCCTACTTTCGGCGTTTCAAGCCATTTTTGGACAATTATATAATCAATTAGAGCTTATAAATCGTCGTT
GATGGTTCCTATTACGACGAGGCACTCGTTTTTTACGCCGAATCGCTCGTAATTACGGACGAGGCAAGAA
GAGACTTGTGTCGGCGAGTTGTCAGGAGCCTCGACGCAGTTCCATTGCTTCAATGTTAAACTAGATTCGC
CCAAGGAGTCTCTTTCTTCGCCCTAGCATACTGTTGCTTCGCCCTAGCATCCTCTTACTTCATCCTAATC
TTCAATTACGCCCAAGCAAATACATTTTACGGTGAAGCAAGTAGCTGTTTCGGCGTTTCAACAGCCTGTT
TGTGTCGGCGAGTTGTGCTCTTTTTTCGCCGAAGTTCCATTGACTCAATGATAAGTAGCCTCGCCCAACC
TGCCTCTAACTTCGCCCAACCTGTCTCTAACTTCGCCCAAGCTGTCTCTAACTTCGCCCAATCATTTTGG
GGGGCTTTCTGGTCTCTTGCATCGTGGTTTCTTCCTTCGCAAAGATTCAACCGGTTGATTCTTTCGTGAA
GCAAGTTCTTGCTTCACCATAAGGTCACGTGATTTCTAGGAACTTTTGGCGAGGAAGTTTAAGAAGCGCC
GTAGACGAAAATAGTGTCGCCGAATTGTTGAATGTAGTTTAAAAACCTCCCTCTTTTCTGGAAAAGCAAG
AAGCGACGCGACAAGAACCCTCTGTTACGTCTCCGTGAGAACTAGCCATAACATGCTGAGGACTCCCGCT
TTCTCCCCCATCCCATCAACTCTCATGAGCATCCAGAACTGTACTTGCAGCTTGAAGTAACTAGCTAGTT
TCTTCAAGCTTACAGGGTGGTATTTTATTGTAGGCCTATCTGTCTATAGATCTTGCTTGTGTGATAGTAA
GTTTTGATCAGCAGTAGTTTTCAGGGGAAAATAGTGGTTGTCTTTTACATCGAGCTCAGTAGAAGATACG
AGTGTTTAAAACATCACGAAATGTTATTCTACCTGTTCACAAAGTTGATTTTTAAAATTTCAAATAATAT
GTAAATGGCACTGCCATGAAATTCGCACAGCATTGTCACAGAAGTACTGCATTATCTTTAACTTTCCATG
CAGAAGGCCAATTTATATTTTTAAATGACTAAGCAAATGGGCATATAGATAGAATAGATAGATAAGATCA
GATCTATAGATAAGATTAGATAGATAGATAGATAGATAGATAGATAGATAGATAGATAGATAGATAGATA
GATAGATAGATTGCACTGACAGTTGTGTAATTCAAATCTGTGTTAAATTTGGATCTATAAATTTGGATCT
ATTATGTTAAATTGTTGATGTTACATAACTGAGATCAATAAAACAATAGCTATTCAGAGATGACTGTTTA
CTATTAGACTGTTACATAATTGTTGATAGTTAATGAAGATTATGTAACCTTAAGTGTTGTTGTGATTTCT
TACTTGGTGAGCCATTTAAGTCATTAACTTTTGTGTTGACTTTTAATTTTCAGTCACTTCTTATTACAAT
CAGGGGTATGGAAATAATTATTTCCAATGTTGATACACATGTTGATTTAAATCCCTGAAATATGTCATTG
GAGGTTGTTATACGACGAGACTGAAATTTTAAAGCTATTTTGGAAGATATTTGAGGTATTTTAATTATTT
ACCAAACTAAACTCTGACTAAACTACTAGCTCCACTTTTATGGCAACTACAGGCACTTCCTGTTTAGGCA
GTCACTGTAGACAGGAGCACTACACAGAATGAGCTGTCCTTGTGCTAAGTGAGCTTCGCTCATGGCTTCC
TTCGGCAGCCATGAGCTGCGCTCATATTATTTTCCTCACATAAAAAGCAAAAAAAATCTAAAATTTTAGG
AGATTTTGGTCACAAAAAGGAGACTTTTGCAAGAAAACTTTTTAGTCAAAAGGAGGTTCTGTCAGAAGGA
GATTTTGTCAAAAGATGATTCTGGGAAAGGCGATTTTTGAAAAAGGAGAGTTTGACTGTCCTCTTGAAAG
GAGTACGCTTTCGTTGTGTGGCTTGTTTTATTCAAAAGAAAGGGTAATATGTTTTTCAATATCTTTCATG
ACTGAGTTGTACACCTACATTGATCAGCACAAGGTCATTGAGACACTTGTTGTCGTAGATCCAGCCGAAG
GCCAACTTCAAAGTCAGTCTTTGCTTGTACATGCACAGACATAAAGGTATCTTTCTATCCTGATAATATT
GCAATGAAATTGATTTCATTTAACTATAGGCCTACGTGCCTGTTGAGAAAGATATTCTGATGAGTAGAAT
GCTTGGAGTCTGCTAGATGAGTTCTTGGAAGATCTTAATGGAATTACATTAGTGGGATCCAGAAGCAAGG
GCAGATAAGAAGAAAAAAGAGACGTGCTTGACAAGGAGGGGGGGGGGGTAATAACACCCCCTCTCTGTTC
ACCCCCCCCCCACCTTCATCCTCCTATCTTCTCTCNATCTTGATGTAAAGGCCTCTCAGCGACTCTCTCA
GGCTTCTGAGTAGAGCTGCAATCTGATTGGCTGTTGGCGCGTGAGCTGCGCTGTGATTGGCCAGTGCCTT
TGAGTAGCCACGCCNCCTGCCTTGCAGCCCCTCAAGAGCTTTCACACTGCTTGATCAGGCTTAATACCAC
GAATCCCACCAAGTGTAGAGCACCCAGCCAACTCCGTGCGATTGTGGACAGGTCGAGGGGGATTGGGCGG
ATGCTGTGAAGTGGGACTTGCAACAGCAACAAGGAAAATCACAACAACAGGACTTGAAACCCAACTTCTT
CTGGAAAACGAGCATCTCTCACATACTCCCAGGTTTAAAACAAGGAGCTCCAGGACAACAGTGTCTGAAT
CAGGTAGGTGATGTTGGATAGCATCCATCTCCAACATCAGCAACAGGCTTCATCAGTAAGCCACATCAGC

AAGCCACATCAGCAGCCTCCACGGCTGAATATCAGTTGGTCCATCAGCTTTTGAGCAGCCTCCTAGCTTG
TGAGAAAATTCTAGGACTTCCTGGCTGCTGTAGACAATCCTTACAACATCCTTCCTGTTCCTGTAGCTTC
CTGTTGAGATGTGAATCTTGCTGATGAATGTGCTGATGTTGTTGATGATGAAGAATTCCTGATGCTTGTG
ATAATTATCGATGTTGTTGTGTCGGTGCTGTTGTCGATATGTCTGTAATTTGTTGATTATATTTTTATCT
TCCGATGATGATGCACATATCCAGCTTCTTTCACCTCCCTGAGATAACATGCAGAATCTTCTCGAAACCT
TCTGAATCACCTTGAATATATACGGGTTTTTTTCTGTTTCATTTCTGAATCCAATGATTTCCTGATTCTT
GTGCTTTATATATTCTCAAATTTATCCGAGAACGCAACCATTTTTCACAGACTCTGCATGGGTGACAGAA
GACATGTTGGAAAGTTCGTTCCATGCACTGGCAGTTAAGCTATGTTCTTTTAGCTTAAAAAGTTGACCCA
TTTGCACAAACTGAGAAGGGTCTATAAAGTGAACATTTTAAGCAGAATTCTTGCAGGCAAAATATCCCTT
CCAAAAAGTACTAGGTAGTAAATATTTTAGGCGACATTATAAATCATACTAGCGGTAATCCAGAAATGTA
ACTTGATTTATGCTTGTAAAATTCTACCATGCAAGTTAGGCCTCTGGAACAGTCCATGATGTAAAAGCAG
GACAAGATTAGGATGTTTTGTACCTGTGATGCTTCAGGTTTCCTCAACTAGATTAGCCTTATAAGCCTTT
TGCAGGTGTTTTTGCTTTTGACATAGACCTAGACCTCCAAATGGAGCTTATATGCCAAACTCGTGAGCAG
TGGGAGGTGCCAGGTCTCTATCTGTCGTTTTGGAGCTTCTCGAATCTTCACAGCCTTGCACTCTTACAAG
TTTTCGAGCTGCTTAACACCACGAAACCAACAAATGCAAATCCACTGGCTCCCTAAGCTTTGAGAGCTCT
CTTATTTGATAGGTCTCTTATTTGAGCTCTTATTCGAGCTCTCCCATTTGCTCCTATTAGAGTTGCTCAC
AAGTCGCACAACTGGGCTTAGAGGAGCTTCAAGGGAACACCCATGACAGACAGTGTTTATAAGGCATCTC
ATCAGCCCCTTCCCTCCAAAAGATTCCCTATAAATATACTTACAATCGAATCAGCCTGAATCAGATTGGC
CTAGCAGCATTTGCCTGCAGCTTTGCCTGCACCTGCCTGCTACTGCTCACTGCAGGAGAGGAGTAAAAGT
TGCCCATCTTCCAACTGCATACATTCAGCATTTTGGCAACACCTACACCACATCTACTTGGATTTTTGTG
AACATATTTCCATTTTGAAATATTTCATCTGACTGAACTTGGACTTGGAAGTCTTGGACATATCAGGAGT
TAGAAATCTGCGCAGGCCAGCATTATGCAGTGAGAATTTGGACACCACTGACCCATCAGCCCAGCTTTGT
GTGCTTCTGGAAGACACTCACACTTGTCTGGCAAACCATGGATCTATTTCTTTTAAATTATCCAGTTCTG
GACTGAAATGTCACAAAGATGGTAAACTAGCTTTCTGACGAGGACTTTGAAAAACTTTTATACACCATTA
CACCTGCAGTCATTAGCTTCAGTTCTTAGTTTTTGACAATGGTGAGCCATTCAGTTTGGCTACAAACTTA
GGAGACTTGGATCACTATTTGATACTTCTGTGTACTGGAACAAGAATTGTATCCTTCTATTGTGATGTCC
AGTTTTGAATATTGGAAACTAACTTCATGTTACCATTTTCAATTTCTAATTCTCACTCGCGCTTGGGATT
TTCGTTTCTGGTATTCTCACATTGATCTTTGACAAGTCGTCTCATATTACCAACACTGGACTTCACCACG
AACCTTTGAGATAAATATTTCACAGAATACTCACTTTTTGAAAATTTTGGTAACCCGCTGAAATTTTGAG
CGGTGTGAGCATCATGTGGAATCAGCTACATCATATTTTTGTCGTCTCTTATATTTTTTTAATGTCTCAA
AATTGATGTTTAAACAATTCGAGACAAATCTATTCTAGTATGAGCCCAGTCATTCGGGCCTAAAGAGCTC
TGAAAATTGAGAAACTTTCAGAAAAACTGTTATTTTGTACGGAAAAACACACATTTGACATTTCTGAATC
GGTAATGCGCAGTTCCCAGAATGTTCCAGAATGTACGCCATTTTGAATTTTGTGACGGGCAGACATTTTG
TCGTTTTGGTAATGCTGCTGGAGTTGATTCTCGTACAGCGAGCAGGTTATCATTGAATGCCAAGGATTTT
TGATGGACTGAATTTTCCCCCTTAGTTTGGATTAAATAAAATAGGTTTTGGAAAAAA*ATG*CTAAACGTTC
GGATGACTGTGGAggcgcgcctt
